# A multi-modal ResNet model to predict coastal fish occurrences using a seascape approach

**DOI:** 10.1101/2025.10.29.682812

**Authors:** Gaétan Morand, David Mouillot

## Abstract

In a context of rapidly declining biodiversity, knowing the distribution of endangered species is critical to ensure the protection of the areas they occupy. To achieve this, species distribution models (SDMs) typically use a range of variables to determine the suitability of an area to a given species, which enables scientists to produce species distribution maps.

Most SDMs are statistically limited in the number of predictors they can take into account, which leads to using summary variables such as temperature yearly average or bathymetry extrema. Deep-learning-based SDMs have been proposed to tackle this limitation by bringing powerful implicit feature extractors. Here, we describe a new model to advance the adaptation of such Deep-SDMs to marine environments and take advantage of the knowledge of the environmental seascape around surveyed points.

We used data from the *Reef Life Survey* data set to predict the presence of 1,796 fish species around all of Australia, in a wide range of climates. The environmental seascape around each point was encoded as a raster image populated with 15 environmental variables, 4 human activity variables, and completed by a 10-year time series of temperature anomaly.

The model performance was highly dependent on the species, with a 0.95 F1 score for the best performing species, *Notolabrus parilus*, but rapidly decreased, with the 100th best species having a 0.38 F1 score.

## 1. Introduction

Species distribution models are a crucial tool for understanding biodiversity, its geographical variations, and the risks related to anthropogenic pressures [1, 2]. Most often, they rely on socio-environmental variables at the surveyed area or observed location and therefore overlook the landscape around these samples.

In the last few years, Deep-SDMs have been used as a way to include the socio-environmental context in the models [3]. In the terrestrial realm, optical satellite imagery was shown to contain a lot of meaningful information, in addition to bioclimatic rasters (average temperature, precipitation, etc.) [4].

In the marine realm, satellite imagery is less significant, as water absorbs light and therefore prevents observation of the underwater environment. Instead, socio-bio-geo-physical variables are used to summarize the local socioeconomic and ecological conditions [5].

## 2. Methods

### 2.1 Occurrence data

In this case study, we used data from the *Reef Life Survey*, which are openly available and provide standardized fish surveys. The data set is strongly skewed towards Australia, which represents 87% of all surveys. We chose to use only these surveys because Australian waters are fully connected and encompass a wide range of socio-environmental conditions (protected areas, tropical to cold temperate climates).

This selection represents 27,437 surveys and 1,796 species, spanning over 103 geographical locations (as defined in the Reef Life Survey metadata). These locations were split with a probability of 60-20-20 percent into train, validation and test data sets, respectively. The split was conducted on the location rather than on the surveys to limit spatial overlap between the three subsets and perform a spatial cross validation.

The *Reef Life Survey* provides abundance data, which were converted to presence-absence for our case study. The average species richness per survey was 19.2 (std=17.8) and the average species prevalence was 1.1% (std=2.9%).

### 2.2 Socio-Environmental variables

For each survey, we downloaded socio-environmental data in a 200km-wide square centered on the survey using the GeoEnrich python package [6]. 19 variables were selected: 15 depicting the natural seascape, and 4 that measure the impact of human activities. These were interpolated to 32 × 32 pixels and stacked into 3-dimensional tensors, as shown in Figure 1. The variables were then normalized into the [0, 1] interval, and outliers (first and last percentiles) were replaced by the corresponding extreme. In addition to this geographical data, we downloaded Degree Heating Week 10-year time series at each survey location. These data were transformed to a 10 × 365 pixel image, subsequently interpolated to 224 × 224 dimensions. The complete list of variables is shown in Table 1.

**Table 1.** 20 layers used as input data. Res. = Resolution. P.S.U. = Practical salinity Unit.

| Variable | Source | Res. | Time Res. | Unit |
| --- | --- | --- | --- | --- |
| FSLEs (strength) | CNES [7] | 0.04° | daily | $days^{-1}$ |
| FSLEs (orientation) | CNES [7] | 0.04° | daily | $degrees$ |
| Bathymetry | GEBCO [8] | 0.0042° | | $m$ |
| Chlorophyll | Copernicus [9] | 4km | daily | $mg.m^{-3}$ |
| SST | Copernicus [10] | 0.05° | daily | $kelvin$ |
| Salinity | Copernicus [11] | 0.083° | weekly | $P.S.U.$ |
| pH | Copernicus [12] | 0.25° | monthly |  |
| Surface Current (u) | Copernicus [13] | 0.25° | daily | $m.s^{-1}$ |
| Surface Current (v) | Copernicus [13] | 0.25° | daily | $m.s^{-1}$ |
| Wind (u) | Copernicus [14] | 0.125° | hourly | $m.s^{-1}$ |
| Wind (v) | Copernicus [14] | 0.125° | hourly | $m.s^{-1}$ |
| Wave Height | Copernicus [15] | 0.2° | daily | $m$ |
| Oxygen | Copernicus [16] | 0.25° | daily | $mmol.m^{-3}$ |
| Mixed layer thickness | Copernicus [11] | 0.083° | daily | $m$ |
| Nitrate | Copernicus [16] | 0.25° | daily | $mmol.m^{-3}$ |
| Protection level | Australian Marine Parks | 0.01° | | $m.s^{-1}$ |
| Distance to port | Global Fishing Watch | 0.01° | | $km$ |
| Population density (log) | Gridded Population of the World | 0.0083° | 5 years | $km^{-2}$ |
| Human gravity | [17] | 0.01° | | $mmol.m^{-3}$ |
| Degree heating Week | NOAA | 0.05° | daily |  |

**Figure 1.**
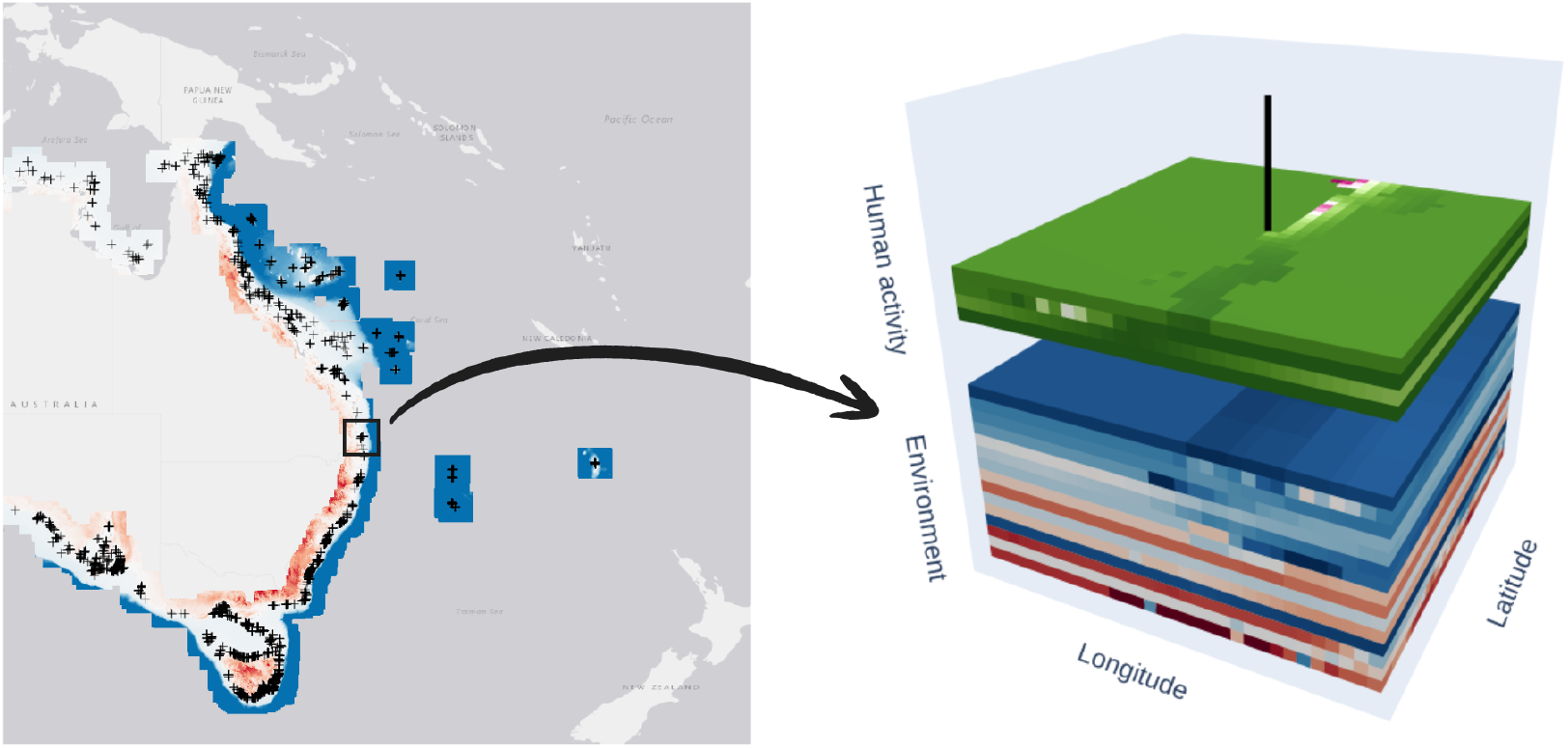
Illustration of the geographic data cube corresponding to a survey. Details on the content of each layer are in Table 1

### 2.3 Model Architecture and training

We used two ResNet18 [18] backbones: one for the environment and human impact layers (19×32×32 pixels inputs) and one for the time series (224 × 224 inputs), as shown in Figure 2. Their outputs were concatenated and normalized with a LayerNorm, then pooled together using a linear layer into a 1,796-long tensor representing predicted probabilities for all species.

**Figure 2.**
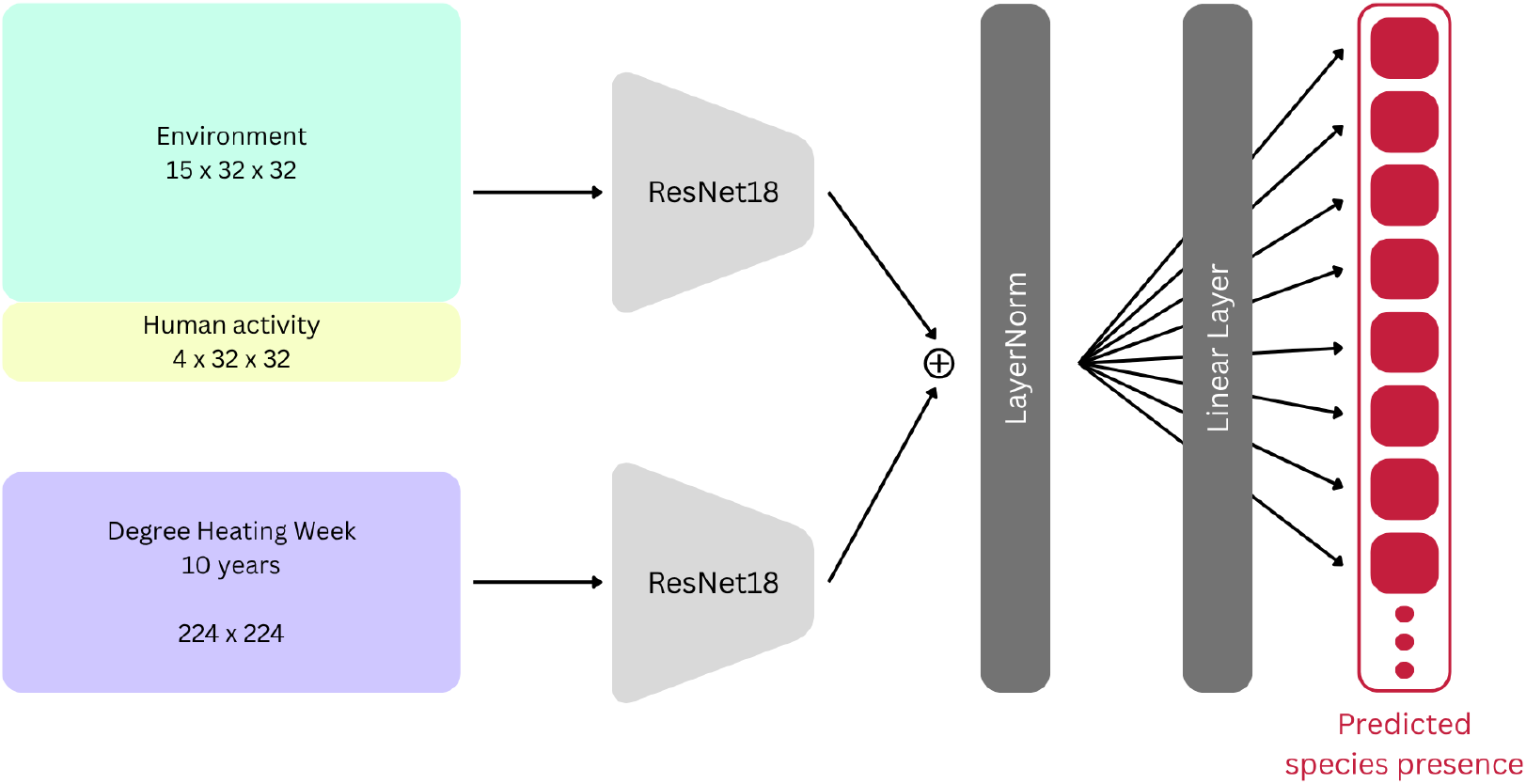
Model architecture

The code, configuration file, and checkpoint are available on Zenodo [19].

No data augmentation was conducted in order to maintain environmental data integrity (*e*.*g*. a northern wind is different from a southern wind).

The model was fit with a cross-entropy loss and a decreasing learning rate until all metrics were stabilized. The asymptotic metrics after 60 epochs were macro-accuracy=0.64 and micro-accuracy=0.99. An important weight decay was used to limit overfitting: 0.5.

### 2.4 Probability cutoff

The threshold for presence/absence has a significant impact on performance metrics. We chose to optimize it on species richness, using the validation data set. We minimized the difference between the mean species richness on the target data and the predicted probabilities, which yielded a 0.235 optimal threshold.

## 3. Results

### 3.1 Species richness

On the test data set, a linear regression of species richness between ground truths and predictions yields a 0.68 R value. The full density plot is shown in figure 3.

**Figure 3.**
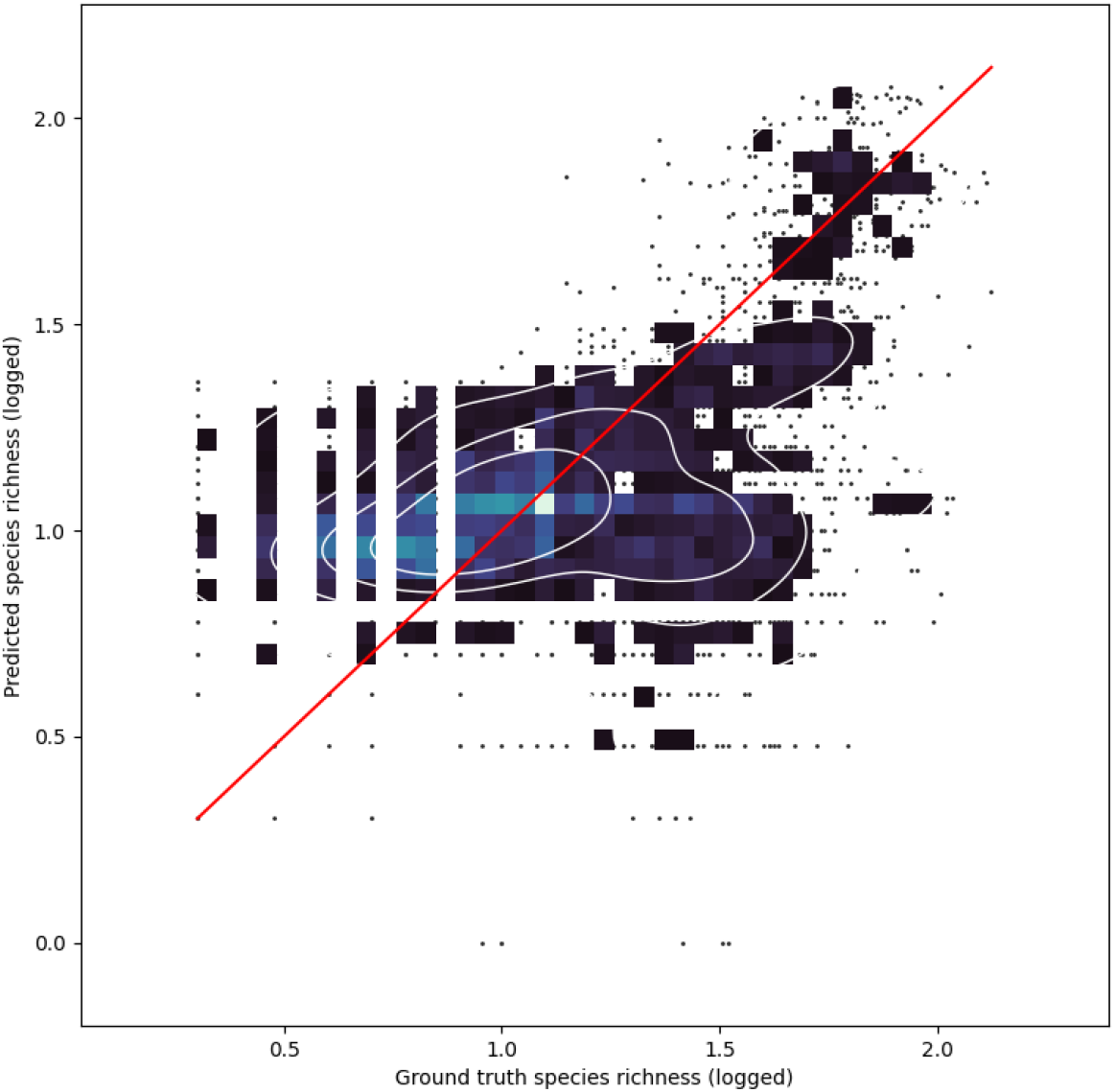
Density plot of ground truth vs predicted species richness on the test data set (N=4,953)

### 3.2 F1 by site

The average F1 score by site, after aggregation of all test surveys, is 0.34 with a 0.17 standard deviation. The map in figure 4 shows the geographical spread of test sites, and the performance for each area.

**Figure 4.**
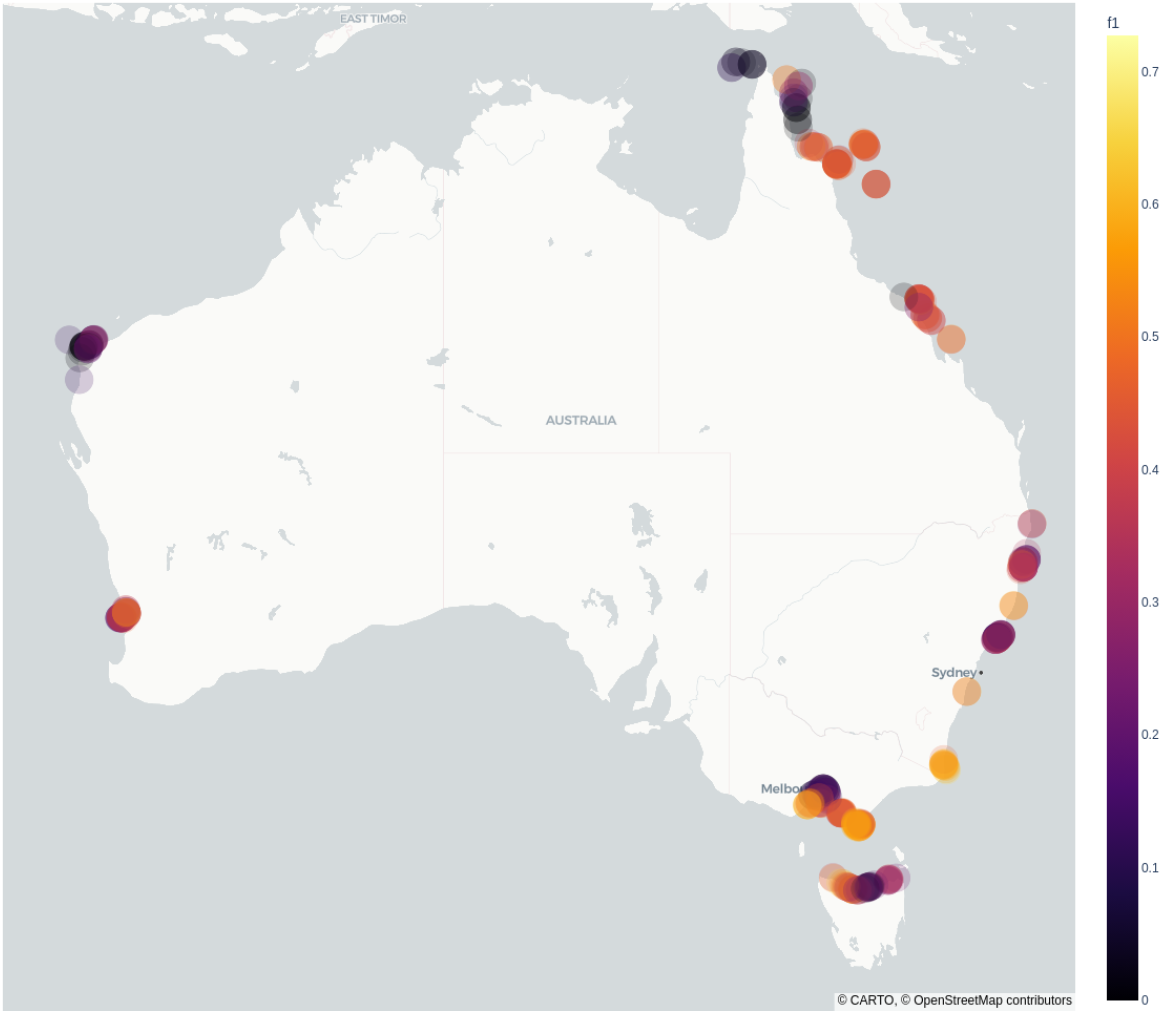
Map of site-wise F1 scores

### 3.3 F1 by species

Species-wise, the F1 score varies significantly. Table 2 shows the best performing species as well as the species ranks corresponding to some F1 thresholds (multiples of 0.1). Figure 5 illustrates the decrease in F1 in relation to species rank.

**Table 2.** Best performing species and associated F1 scores.

| Rank | Species | F1 |
| --- | --- | --- |
| 1 | *Notolabrus parilus* | 0.950 |
| 2 | *Notolabrus tetricus* | 0.866 |
| 3 | *Coris auricularis* | 0.785 |
| 4 | *Caesioperca rasor* | 0.780 |
| 5 | *Acanthochromis polyacanthus* | 0.763 |
| 6 | *Halichoeres melanurus* | 0.761 |
| 7 | *Pomacentrus moluccensis* | 0.758 |
| 8 | *Pomacentrus wardi* | 0.738 |
| 9 | *Chrysiptera rollandi* | 0.728 |
| 10 | *Chelmonops curiosus* | 0.706 |
| 23 | *Coris batuensis* | 0.600 |
| 55 | *Paracirrhites arcatus* | 0.511 |
| 89 | *Pomacentrus coelestis* | 0.400 |
| 135 | *Halichoeres trimaculatus* | 0.303 |
| 182 | *Scolopsis monogramma* | 0.200 |
| 251 | *Trygonoptera imitata* | 0.100 |

**Figure 5.**
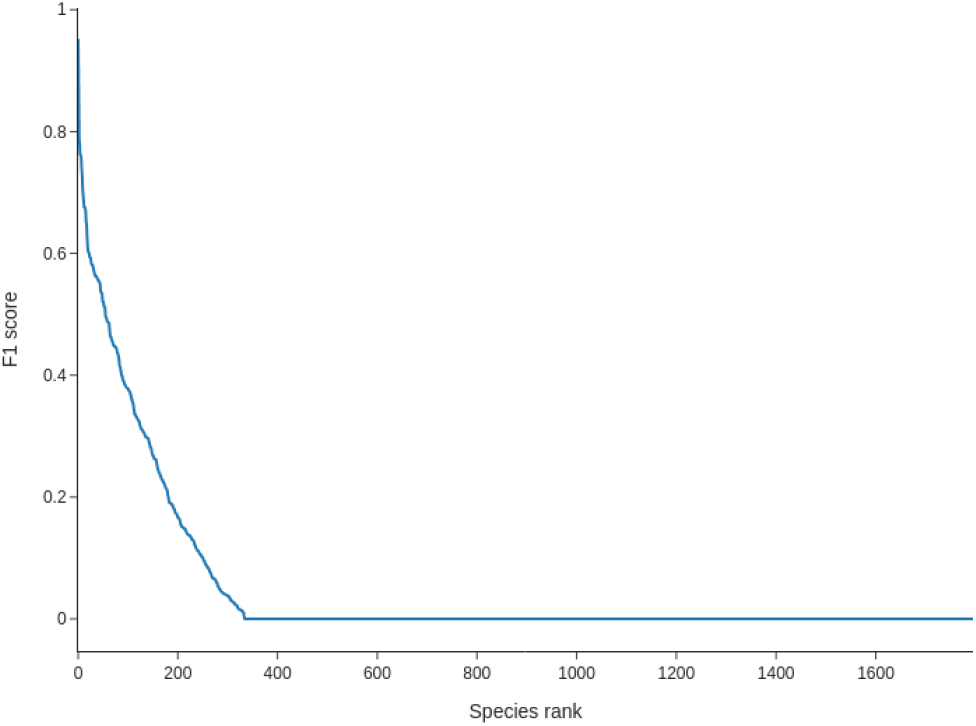
F1 score by species

## 4. Conclusion

The model was able to learn the association between socio-environmental rasters and occurrences for some species but not others. Although performance was highly dependent on the species and the location, there was a significant success overall in predicting species richness. Such models and approaches are first critical steps towards better predictions of biodiversity patterns across space and time that are urgently needed by managers, scientists and organizations.

## 5. Acknowledgment

This work has been mainly funded by the IA-Biodiv ANR programme FISH-PREDICT project (ANR-21-AAFI-0001-01).

This project was provided with computing AI and storage resources by GENCI at IDRIS thanks to the grant 2025-AD011013891R2 on the supercomputer Jean Zay’s V100 partition.

